# Postembryonic Development and Morphology of the Reproductive System in the Acoel *Hofstenia atroviridis*

**DOI:** 10.64898/2025.12.15.692464

**Authors:** Rupandey Parekh, Coline Hermine, Yohei Otomo, Toru Miura

## Abstract

Reconstructing ancestral reproductive systems is essential for understanding the evolution of bilaterian body plans, yet the origin and development of reproductive organs remain poorly characterized. Here, we investigate postembryonic sexual development in the xenacoelomorph *Hofstenia atroviridis*, a basal acoel species, to gain insight into early bilaterian reproductive evolution. Individuals were reared from eggs to adulthood, and the ontogeny of reproductive organs was examined using histology and immunohistochemistry with muscle and neural markers. *H. atroviridis* is a protandrous simultaneous hermaphrodite, with sexual maturity correlated with body size rather than age. The male copulatory system comprises a seminal vesicle, granular vesicle, penis with a penile bulb, and a single copulatory stylet, accompanied by regionally specialized musculature and innervation. In contrast, the female reproductive system consists of paired, asaccate ovaries containing large, follicle cell-bound oocytes and lacks a discrete gonopore or copulatory organ. Fertilization occurs via traumatic insemination, and eggs are likely released through the mouth. Despite the organism’s overall morphological simplicity, the male reproductive system exhibits pronounced structural differentiation. These findings suggest that sexual selection acting on a hermaphroditic ancestor may have contributed to the early diversification of bilaterian copulatory organs and establish *H. atroviridis* as a useful model for studying the evolutionary origins of animal reproductive systems.

## Introduction

One of the essential contributions of evolutionary developmental biology (evo-devo) to reconstruct the ancestral form of life has been clarifying the developmental processes that shaped the evolution of morphological traits in basal species of the metazoan phylogeny (Hejnol and Martindale, 2008). Considerable progress has been made in reconstructing several features of the last common bilaterian ancestor (LCBA), also known as the Urbilaterian, including axial polarity (Genikhovich and Technau, 2017), body regionalization (Moreno et al., 2011; Luo et al., 2018), appendages (Tarazona et al., 2019), segmentation (Balavoine and Adoutte, 2003), photosensitive cells (Nilsson, 2021), circulatory systems (Hartenstein and Mandal, 2006; Monahan-Earley et al., 2013; Stephenson et al., 2017), gut (Nielsen et al., 2018), and nervous systems (Furness and Stebbing, 2018; Martín-Durán and Hejnol, 2021). In contrast, the reproductive system remains far less explored, although the reproductive system of the Urbilaterian must have been highly efficient, given the evolutionary success of the bilaterian clade (Extavour, 2007). While subsequent work has begun to address this gap (Sasson and Ryan, 2017; Leonard, 2018), our understanding of early bilaterian reproductive systems remains incomplete.

Basal taxa are critically important for reconstructing ancestral traits at the base of major clades. For Bilateria, a key group in this context is the phylum Xenacoelomorpha, though its exact phylogenetic placement remains debated. While the prevailing view places Xenacoelomorpha as sister to Nephrozoa (Protostomia + Deuterostomia) (Hejnol et al., 2009; Paps et al., 2009a, b; Cannon et al., 2016; Rouse et al., 2016), other studies support a sister relationship with Ambulacraria within a monophyletic or paraphyletic Deuterostomia (Philippe et al., 2011, 2019; Kapli and Telford, 2020; Kapli et al., 2021). The most recent and largest phylogenomic analysis of bilaterians, supports Xenacoelomorpha as a sister to all other bilaterians (Álvarez-Presas et al., 2024).

Acoela, an order within Xenacoelomorpha comprising approximately 450 species, are small, soft-bodied marine worms with highly simplified body plans and rapidly evolving genomes (Duruz et al., 2020; Atherton and Jondelius, 2022; Martinez et al., 2024). They lack a coelom, a through gut, and circulatory, excretory, and respiratory systems, as well as larval stages and basal laminae. They also lack ciliary and rhabdomeric eyes. Despite this overall simplicity, the few organs they possess exhibit considerable variation in structural organization (Jondelius et al., 2011; Achatz et al., 2012; Brusca et al., 2016; Asai et al., 2022; Hookabe et al., 2024). Among these, the reproductive system is particularly diverse, yet current knowledge is limited to only a few acoel lineages (Jondelius et al., 2011). Therefore, reconstructing ancestral reproductive systems remains challenging — not only in acoels (Jondelius et al., 2011; Abalde and Jondelius, 2025), but also across bilaterians more broadly.

Furthermore, research on acoel reproductive systems is hampered by significant sampling biases — geographically toward the Scandinavian coast and North Atlantic (Jondelius et al., 2019), and taxonomically toward Crucimusculata, the more derived and faster-evolving of the two major infraorders (Jondelius et al., 2011; Abalde and Jondelius, 2025). Of the ∼450 described acoel species, only a small fraction have detailed anatomical descriptions of reproductive systems, and only two studies to date have investigated sexual development (Kozloff, 1965; Chandra et al., 2025). While morphological descriptions of adult reproductive systems are available for many species, studies of reproductive system development are exceedingly rare. This is not a problem unique to acoels, a similar gap in developmental data for reproductive systems across Bilateria have been noted (e.g., Extavour, 2007; Leonard, 2018).

To address this gap, we investigated sexual development in *Hofstenia atroviridis* Bock, 1923, a member of infraorder Prosopharyngida — a simpler and more basal clade than the derived Crucimusculata (Jondelius et al., 2011; Achatz et al., 2012; Martinez et al., 2024). Unlike its better-studied relative *Hofstenia miamia* Correa, 1960, *H. atroviridis* shows a uniform dark brown coloration throughout its life cycle. In coastal areas of Japan, this species can be collected in the field and maintained in the laboratory with relative ease (**Fig. 1**). Here, we described the postembryonic development of its reproductive system and characterize the anatomy of the mature organs. *H. atroviridis* has a complex anterior male copulatory organ and posterior oocytes as the female system. This study clarified that this species was a protandrous simultaneous hermaphrodite, with sexual maturity determined more by body size than by age, as also reported in *H. miamia* (Chandra et al. 2025). This allowed us to divide its postembryonic development into five size-based stages. Using commonly available antibodies for neural markers, we identified clear regionalization in and around the male copulatory organ. We hypothesize that sexual selection acting on a hermaphroditic ancestral system may have driven the divergence between the morphologically simple female system and the more elaborate male genital structures.

**Figure 1.**
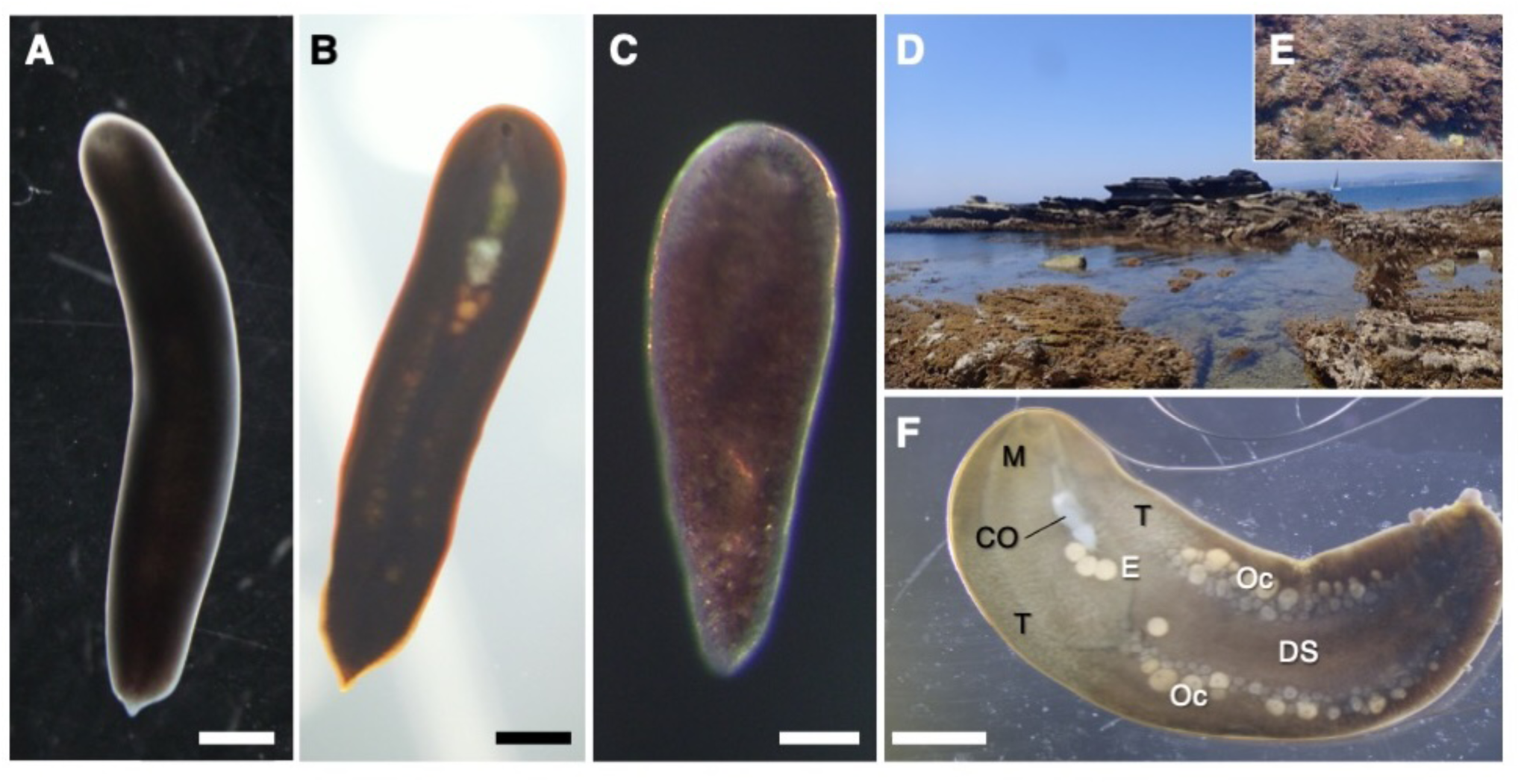
Overview of the acoel worm, *Hofstenia atroviridis*. (A, B) Dorsal (A) and ventral (B) view of an adult individual. (C) Dorsal view of a hatchling in 24 hours old. (D) Collection site of *H. atroviridis* in this study. *H. atroviridis* inhabits in shallow waters of rocky shores, even in tide pools. (E) Animals are usually found among coralline red algae (Rhodophyta sp.). (F) Ventral view of a larger individual (≥ 8.0 mm in body length). Rough structure and positions of internal organs, mainly reproductive systems, are visible. Scale bars: (A, B, F) 1 mm, (C) 200 μm. Abbreviations: CO, male copulatory organ; DS, digestive E, egg; M, mouth; Oc, oocytes; T, testis.

## Materials and Methods

### Animal Husbandry

To obtain wild populations of *Hofstenia atroviridis*, *Corallina* sp. algae were collected from tide pools at Araihama Beach in Misaki, Kanagawa, Japan (35°09’33”N, 139°36’42”E). These pools were regularly replenished by wave action and characterized by sandy, not muddy, sediment. The algae were washed by hand to collect *Hofstenia* individuals, which were then transported to the laboratory. Worms were maintained in filtered seawater (FSW) in plastic containers (84 x 57 x 44 mm; Entec, Niigata, Japan) with perforated lids for oxygen exchange. To induce egg production, animals were kept in an incubator at 25°C under a 12 h light/dark cycle, mimicking the condition of breeding season (July to mid-September). Containers were cleaned, seawater was replaced, and worms were fed cod roe three times per week.

When eggs were found in the rearing containers, they were gently scraped from the container surfaces using a pipette and transferred to 6-well plates (Costar 6-well Clear TC-treated Multiple Well Plates, Corning Inc., Corning, NY, USA). Water in the wells was changed three times per week. Upon hatching, F1 hatchlings were collected with a pipette and maintained under the same conditions as adults, although they were fed a suspension of crushed artificial fish food (Megabyte Red, Kyorin Co., Ltd., Hyogo, Japan). Since *H. atroviridis* is cannibalistic, individuals that visibly outgrew their tankmates were separated. Precise measurement of juveniles was not feasible for routine maintenance; instead, individuals were visually sorted by body size, unless used for experiments.

### Measurement of Body Size

Body length was a key variable in this study. Animals were first anesthetized in 3.5% MgCl₂ dissolved in distilled water and saturated menthol dissolved in FSW, then transferred to an embryo dish containing 1-2 drops of the same anesthetic solution. They were examined under a stereomicroscope (SZX16, Olympus Corp., Tokyo, Japan), and images were captured using a mounted camera (BV26, Olympus Corp., Tokyo, Japan) and cellSens Standard software (Olympus Corp., Tokyo, Japan). The polyline measurement tool implemented in the software, was used to trace the midline of each specimen, and the resulting length was recorded.

### Histological Observations

#### Paraffin Sections

Animals were anesthetized and measured as described above, then fixed in FAA (formalin: ethanol: acetic acid = 6: 16: 1) for 3 hours on a shaker. Samples were subsequently transferred to 70% ethanol at room temperature (RT) overnight, or for at least 2 hours. For juveniles too small to handle directly in paraffin, samples were stained overnight in 70% ethanol containing a few drops of eosin. For easier manipulation, specimens were embedded in a thin layer of 0.1% agarose (prepared by mixing 1.5 mL PBS of 1% agarose in an embryo dish), washed twice in PBS, and allowed to solidify. The agarose blocks containing the samples were trimmed with a scalpel and subjected to dehydration.

Dehydration was performed on a shaker at RT for 20 minutes each in the following order: 90%, 95% (twice), 100% (twice) ethanol. Samples were then washed twice for 5 minutes each in xylene, followed by 30 minutes in a 1:1 xylene/paraffin mixture. This was followed by three 30-minute incubations in paraffin at 60°C. Sections were cut using a rotary microtome (No. 820, Spencer, Buffalo, USA), mounted on glass slides, and dried overnight on a slide warmer at 37°C (PS-53, Tissue-Tek (Sakura Finetek), Tokyo, Japan). Slides were stained with hematoxylin and eosin and mounted in Malinol (Muto Pure Chemicals, Tokyo, Japan), then observed under an optical microscope (BX51, Olympus Corp., Tokyo, Japan) and images were captured using a mounted camera (DP47, Olympus Corp., Tokyo, Japan) and cellSens Standard software.

#### Resin Sections

For juveniles ∼2 mm or smaller, paraffin embedding proved unreliable due to brittleness of the agarose block during slicing. Instead, resin sections were used. Specimens were anesthetized, measured, and embedded in agarose as above, but the agarose block was cut using a stencil shaped to match the tapered end of a 1.5 mL PCR tube. Samples were fixed in FAA for 3 hours and soaked overnight in 70% ethanol with eosin. The next day, they were rehydrated in 50% and 25% ethanol (20 minutes each), followed by overnight incubation in distilled water. Dehydration was then carried out sequentially in 25%, 50%, 75%, and 100% ethanol (20 minutes each), followed by three 30-minute washes in 100% acetone. Samples were then embedded in Technovit 8100 resin (Kulzer GmbH, Hanau, Germany) within 1.5 mL PCR tubes following the manufacturer’s protocol. The resin was allowed to cure at RT for 3–4 days. Once hardened, resin blocks were removed from the tubes, sectioned using a microtome, mounted on slides in distilled water, and air-dried overnight. Slides were stained with hematoxylin and eosin, mounted in Malinol, and observed under an Olympus BX51 optical microscope.

### Immunohistochemistry

Whole-mount immunostaining was performed following a modified protocol from Hulett et al. (2020). Animals were anesthetized, measured, and fixed in 4% paraformaldehyde in filtered seawater (FSW) for 30 min with agitation. Samples were washed in 0.1% Triton X-100 in PBS (PBT), permeabilized in 100% methanol for 10 min, rewashed, and blocked in 10% sheep serum in PBT for 30 min. Primary antibodies against serotonin (5-HT; 1:500, Immunostar) and tyrosinated tubulin (1:300, Sigma-Aldrich) were diluted in blocking solution and incubated for 72 h at 4 °C. After washing in PBT, samples were incubated for 24 h at 4 °C with Alexa Fluor 488-conjugated anti-mouse secondary antibody (1:200, Invitrogen) diluted in BBT/NGS. Nuclei were counterstained with DAPI (1 µL/mL PBT) for 30 min, followed by rinses in PBT. F-actin was visualized using Alexa Fluor 488– or iFluor-647–conjugated phalloidin (both 1:500; Cell Signaling Technology and Abcam, respectively) following Srivastava et al. (2014).

Due to high background in whole mounts, smooth muscle actin immunostaining was performed on paraffin sections. Samples were fixed, embedded, sectioned, and mounted as described above for paraffin histology. Sections were deparaffinized in xylene, rehydrated through an ethanol series, briefly stained with eosin, and washed in PBT. Blocking was performed in BBT followed by BBT/NGS. Sections were incubated overnight at 4 °C with anti-smooth muscle actin antibody (GT445; 1:500, GeneTex), washed, re-blocked, and incubated with Alexa Fluor 488-conjugated anti-mouse secondary antibody for 2 h at room temperature. After final washes, sections were prepared for mounting.

All samples were mounted in VECTASHIELD (Vector Laboratories) and imaged using a confocal microscope (FV3000, Olympus). Images were processed with FV31S-SW software (Olympus).

## Results

### Overview of the Study Species

The focal species *Hofstenia atroviridis* (**Fig. 1**) is commonly found in association with coralline algae (*Corallina* sp.: Rhodophyta) (**Fig. 1D, E**) in subtidal areas and tide pools at least knee-deep, where regular wave action ensures clean, oxygen-rich water. While the dorsal surface is featureless (**Fig. 1A**), the ventral side of larger individuals (7–8 mm) clearly reveals rough structures and positions of both male and female reproductive systems, in addition to the mouth and digestive syncytium (**Fig. 1B, F**). The mouth was opened at the anterior tip of the body, followed by male copulatory organs and eggs. A pair of testes (the areas of spermatogenesis) is observed on both sides of the pharynx, although there is no obvious boundary to the surrounding tissues (**Fig. 2A**). The digestive syncytium is located along the midline of the posterior half of the body. Paired ovaries containing various sizes of oocytes are located in the longitudinal area stretching from the anterior edge of the digestive syncytium to a few millimeters short of the posterior end. Wild-caught adults exhibited seasonal variation in size and abundance. From October to May, worm numbers and sizes remain low, averaging 4–5 mm in length. As temperatures rise in June, both abundance and size increase, peaking between July and September, which appears to correspond to the breeding season. Many individuals collected during this period exceeded 6 mm in length.

**Figure 2.**
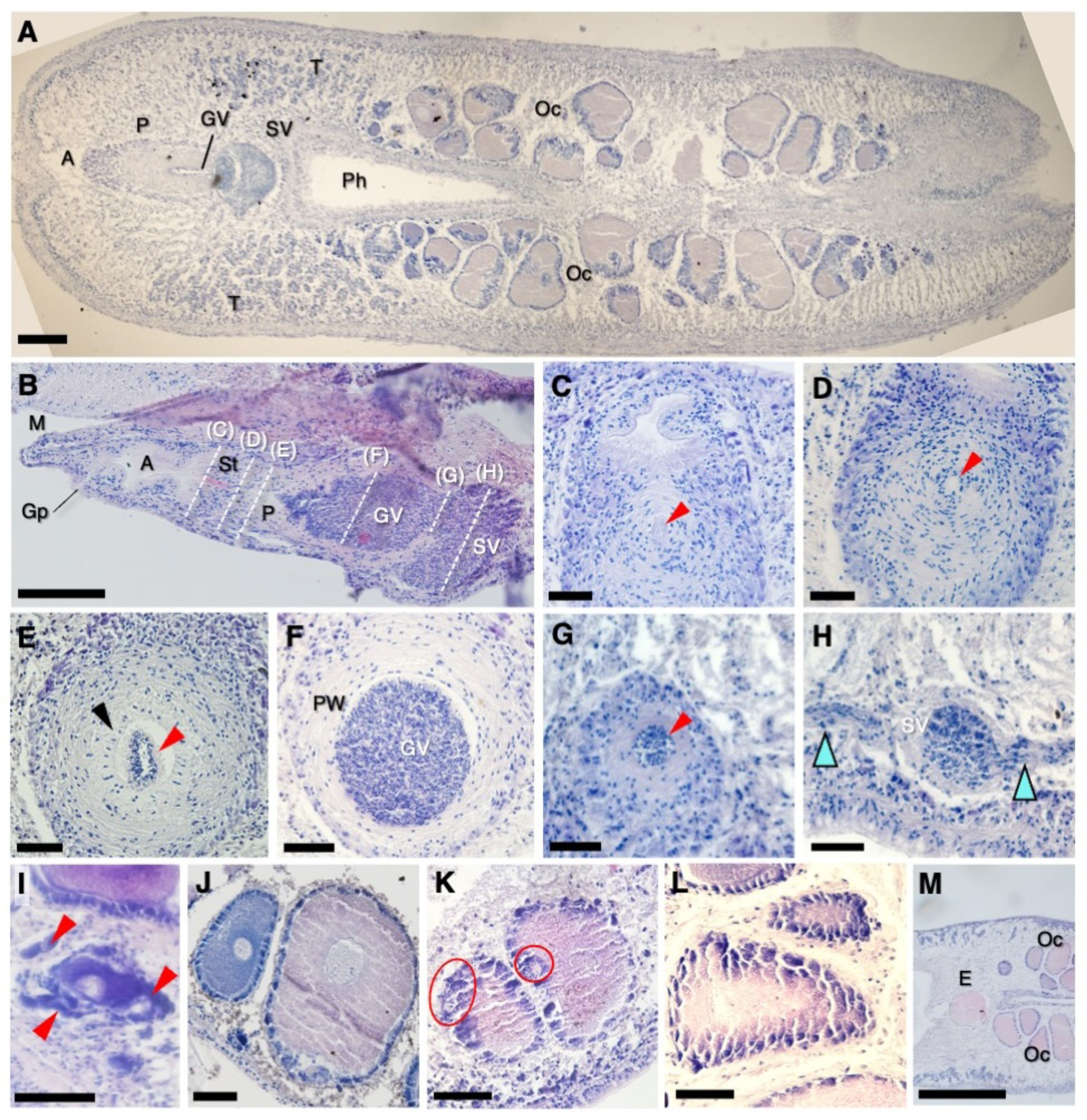
Histological sections of an adult individual of *Hofstenia atroviridis*. Anterior is to the left in horizontal (A, B, E-H) and sagittal sections (C, D), and dorsal is up in transverse sections (C, D). (A) Horizontal section through the whole body, containing both male and female reproductive organs. (B) Sagittal section of the male copulatory organ of an adult. Dashed lines indicate positions of transverse sections. (C–H) Transverse sections of the male copulatory organ, from anterior to posterior. (C) Antrum and anteriormost penis. Arrowhead: stylet. (D) Penis, cut at level of stylet tube (arrowhead). (E) Penis, cut at level of anterior GV (red arrowhead). Black arrowhead: columnar epithelium of penis. (F) Granular vesicle surrounded by penial wall. Note the absence of columnar epithelium. (G) Aperture at the base of germinal vesicle. (H) Seminal vesicle with “channels” (elongated testes follicles) joining from both sides (right blue arrowheads). (I–M) Horizontal sections showing oocytes. (I) Nascemt oocyte located near the body wall. (J) A slightly older oocyte surrounded by precursors of follicle cells (arrowheads) (K) A pre-vitellogenic (left, blue) and vitellogenic (right, pink) oocyte surrounded by a layer of follicle cells. (L) Vitellogenic oocytes at the time of fertilization. Red circles indicate allosperm. (M) Horizontal section showing a pre-released mature egg just posterior to the pharynx. Scale bars: (M) 500 µm, (A) 200 μm, (B) 100 µm, (C-L) 50 μm. Abbreviations: A, antrum; E, egg; Gd, gonoduct; Gp, gonopore, GV, granular vesicle; M, mouth; Oc, oocytes; Pw, penis wall; Ph, pharynx; S, stylet; SV, seminal vesicle; T, testes.

### Structure of the Mature Male Copulatory Organ

Histological sections revealed the structure of reproductive organs in greater detail than surface views, especially in case of the male organs (**Fig. 2A**). Testes were assacate and follicular (**Fig. 2A, B, Supplementary Figure S1, S2**). The copulatory apparatus consisted of the seminal vesicle, granular vesicle, penis, an intromittent stylet, the antrum masculinum and the gonopore, opening just posterior to the mouth (**Fig. 2B–H**). The seminal vesicle was usually rounded, and was enclosed by a muscular wall. Though the seminal vesicle was typically smaller than the granular vesicle (**Fig. 2B**) at its maximum capacity, it could be equal to the volume of the granular vesicle and penis combined. There seemed to be no openings in the wall of the seminal vesicle for sperm entry from testes. Instead, there were follicles, elongated into channels, suggesting that sperm simply pushed right through the wall to enter the vesicle (**Fig. 2C**). Only mature sperm were found in the seminal vesicle (**Fig. 2D–H**). The granular vesicle occupied a large part of the copulatory organ; it was fed by the seminal vesicle through an aperture in its bottom and was surrounded by the penis (**Fig. 2D, E**). An intromittent stylet was observed at the anterior tip of the penis, projecting into or just behind the antrum (**Fig. 2B**).

### Structure of the Mature Female Organ

The adult female reproductive system of *Hofstenia atroviridis* was comprised of paired ovaries flanking the digestive syncytium in the ventral half of the body, and a relatively dorsal group of mature eggs located just posterior to the pharynx (**Fig. 2A**). Consistent with ancestral acoel gonads being asaccate (Jondelius et al., 2011; Achatz et al., 2012), the ovaries were unbound by membranes and are not divided into germaria and vitellaria. Oocytes were characteristically huge and numerous. There was no antero-posterior arrangement of oocytes according to maturity, suggesting that oogonia were not restricted to either end of the ovaries, but randomly appeared throughout ovaries. In the ovaries, oocytes of all stages had follicle cells (except the most nascent ones) and giant germinal vesicles (**Fig. 2I–L**). The youngest oocytes, easily discernible by their giant nuclei and vivid blue hue in HE sections, were found close to the ventral body wall. Soon after, they became surrounded by follicle cell progenitors, identified as such because they were larger than the follicle cells around older oocytes and were of an irregular and flattened shape (**Fig. 2I**). As the oocyte matured, it grew larger and more ellipsoid. Simultaneously, follicle cell progenitors proliferated to form a layer of columnar follicle cells (helper cells) completely encasing the oocyte. Oocytes with ongoing vitellogenesis had violet to pink hues in HE sections due to partially increased affinity for eosin, since vitellogenin was acidophilic (**Fig. 2J–L**). The follicle cells adjacent to the oocyte were columnar with their bottom halves swollen with vitellogenin granules; outer layers are more cuboidal (**Fig. 2L**). These granules are incorporated into the oocyte cytoplasm either by follicle cells actively releasing granules or by incorporation of the follicle cells themselves. Once vitellogenesis was complete, follicle cells took on a flattened shape (**Fig. 2J**). Oocytes at this stage were stained pink in HE sections and were called vitellogenic oocytes. *H. atroviridis* was found to lack female copulatory organs and a female gonopore, as observed in *H. miamia* (Chandra et al., 2025). Oviposition likely occurred through the mouth (**Fig. 2M**).

### Detailed structure of the copulatory organ

Details of the structurally complex male reproductive system, particularly the male copulatory organ, were revealed primarily by immunohistochemistry (**Fig. 3A, B, Supplementary Figure S3**). Testes contained spermatogonia, spermatocytes and spermatids, distinguishable from the shape (**Fig. 3C–E, Supplementary Figure S1**). Notably, spermatids had extremely elongated nuclei, as observed in other acoels (**Fig. 3F–H**; Boyer and Smith, 1982). Spermatocytes arose close to the anterior-lateral body wall, moving ventromedially towards the copulatory organ as spermatogenesis progressed. The follicles did not show distinct zonation, but ongoing spermatogenesis during their ventromedial progression did result in a tendency for more advanced stages to occur more medially (**Fig. 3D, E**). Organization into follicles appeared to be caused by passage through the testes of parenchymal muscles linking the pharynx to the body wall. To some extent, parenchymal muscle funneled developing sperm in the required direction. Actual ducts appeared to be absent.

**Figure 3.**
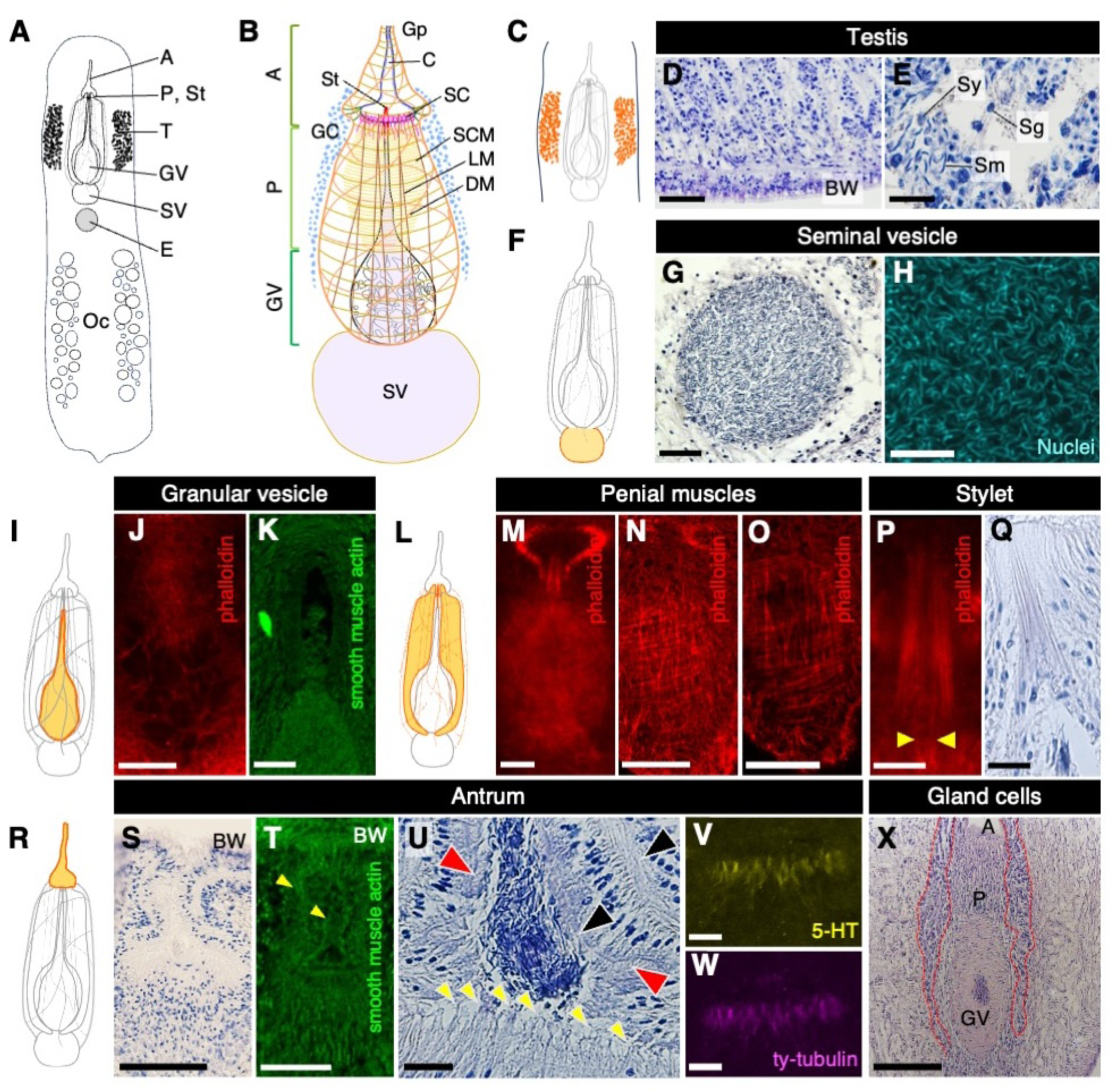
Detailed morphology and components of the adult male reproductive system. Anterior is to the top, unless mentioned. (A-B) Schematic diagram of reproductive organs in whole body (A) and detailed morphology of male reproductive system (B) of an adult individual. (C-E) Testis. (C) Diagram of testes on both sides. (D) Horizontal section of the left testis. Anterior is to the left. (E) Magnified view of the horizontal section of the testis, containing every stage of male germ cells. Anterior is to the left. (F-H) Seminal vesicle. (F) Diagram of the seminal vesicle. (G) Horizontal section of the seminal vesicle. (H) Nuclei of mature sperm inside the seminal vesicle stained by DAPI. (I-K) Granular vesicle. (I) Diagram of granular vesicle. (J, K) Fluorescent images of granular vesicle showing reticulation of its lining (J) and its general shape (K). (L–Q) Penis and stylet with F-actin labeled by phalloidin. (L) Diagram of penis and stylet. (M) Dense layer of penial circular muscles, (N) layer of basket-like muscle external to circular muscles, (O) layer of straight, thick muscles internal to circular muscles. (P) Magnified view of the stylet. Yellow arrows indicate slender actin tube inserting into stylet. (Q) The penile bulb, a layer of hardened tissue surrounding the stylet. (R-W) Antrum. (R) Diagram of the antrum. (S) Horizontal section. The antrum wall is continuous with the body wall (T) Labeling with smooth muscle actin highlights the antrum’s elongated conical shape. The penile bulb is visible inside the antrum’s bulbuous posterior end. (U) Horizontal section. Double wall of the antrum, cilia and roots of hair-like sensory projections are indicated by black, red and yellow arrowheads, respectively. (V, W) Fluorescent images of hair-like sensory projections, stained with 5-HT (V) and tyrosinated-tubulin antibody (W). (X) Horizontal section showing gland cells surrounding the copulatory organ from the posterior part of the antrum to the penis and granular vesicle (red dotted line). Scale bars: (X) 500 μm, (D, N, O, S) 100 μm (G, J, K, M, T) 50 μm, (E, H, P, Q, U) 20 μm, (V, W) 15 μm. Abbreviations: A, antrum; BW, body wall; C, cilia; cm, circular muscles; E, egg; DM, diagonal muscle; GC, gland cells; Gp, gonopore; GV, granular vesicle; LM, longitudinal muscle; P, penis; Ph, pharynx; SC, sensory cells; Sg, spermatogonia; St, stylet; SV, seminal vesicle; Sy, spermatocyte; T, testis.

The granular vesicle was teardrop-shaped with a bulbuous posterior and elongated anterior running up to the copulatory stylet (**Fig. 3I–Q**). The rounded posterior half of the granular vesicle was lined by a reticulated layer of cells (**Fig. 3J**) and was surrounded by the thick muscular wall of the penis (**Fig. 2F**). The anterior granular vesicle took the form of a canal passing through the penis lumen itself (**Fig. 3K**). The bulk of the penis was made of tightly arranged circular muscle fibers (penial circular muscle; **Fig. 3M**) that were contiguous with those surrounding the granular vesicle (**Fig. 2B**). External to the circular muscles were diagonal muscles woven into a thin basket-like layer (**Fig. 3N**), and internal to them was a layer of thick, longitudinal muscle fibers (**Fig. 3O**). The anterior end of the penis protruded slightly into the basal chamber of the antrum and housed an intromittent stylet (**Fig. 3P, Q**). This penial tissue around the stylet was of a more hyaline texture in histological sections compared to the fibrous texture of the rest of the penis, indicating some form of hardening. The stylet was composed of actin needles and sat atop a very slender and straight tube (**Fig. 3P**).

Anterior to the penis was the antrum masculinum, so called because it connects only the male reproductive system to the external environment (henceforth, just “antrum”) (**Fig. 3R, T**). From the male gonopore just posterior to the mouth, it extended posteriorly as a conical passage which flared at its base into a small cavity (**Fig. 3U**). Its wall was bilayered and continuous with the two layers of the body wall (**Fig. 3S, T**). The antral lumen had ciliated walls and the base was ringed with posterior-facing and hairlike projections (**Fig. 3U-W**). These projections appeared to extend from the base of the outer wall. As those were stained by both 5-HT and tyrosinated tubulin (**Fig. 3V, W**), they might have a mechanosensory function, as suggested by Chandra et al. (2025). In addition, numerous gland cells were densely distributed around the male copulatory organ, roughly from the posterior part of the antrum to the anterior part of the granular vesicle, as evidenced by a HE-section (**Fig. 3X**).

### Stages of Sexual Development

It was hypothesized that body size is a key factor influencing the developmental stage of the reproductive system. Therefore, individuals were categorized by body length as follows: 0.5–0.7 mm (hatchlings), 1.4–3.0 mm (immature males), 3.0–5.0 mm (immature males and females), 5.0–6.0 mm (mature males but immature females), and ≥6.0 mm (mature males and females). To examine reproductive development, we conducted detailed observations and descriptions of the histological characteristics (**Fig. 4**).

**Figure 4.**
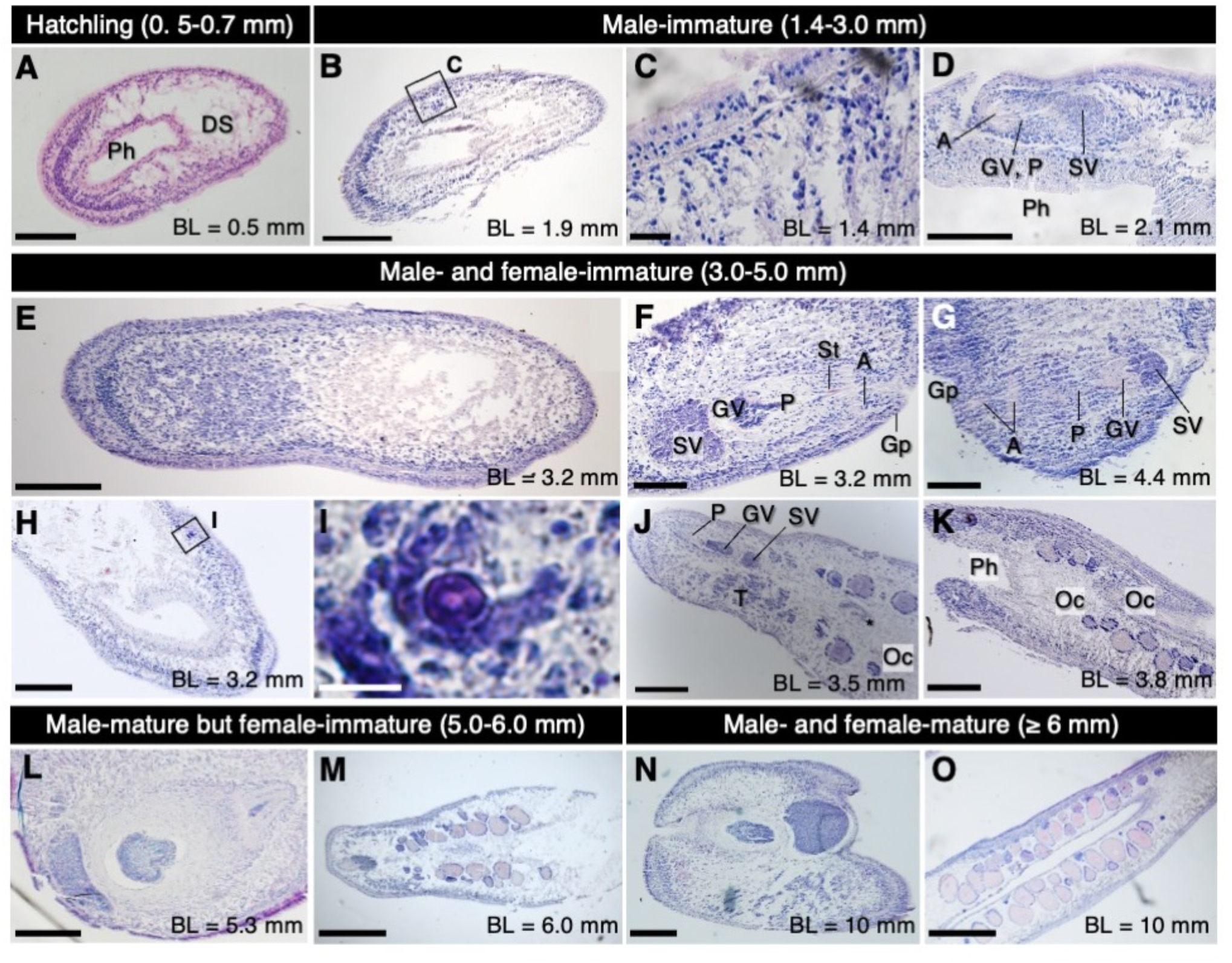
Reproductive development in *Hofstenia atroviridis* is protandrous and correlated to body size. Horizontal sections unless mentioned. (A) Hatchling without any developing gonads, with 0.5-0.7 mm body length (BL). (B–D) Male immature individuals, with 1.7–3.0 mm BL. (B) Spermatogenesis starts at testes. The black-framed box indicates the magnified area showed in (C). (C) Magnified view of the left testis of (B). (D) Rudimentary copulatory organ. (E–K) Male- and female-immature individuals, 3.0–5.0 mm in BL. (E) Whole body. (F, G) Sagittal (F) and diagonal section (G) of an immature male reproductive organ. (H) Posterior half of body in where oocytes start developing. The black-framed box indicates the magnified area showed in (I). (I) Magnified view of a developing oocyte. (J) Individual with a mostly differentiated copulatory organ and a small number of mostly pre-vitellogenic oocytes, 3.5 mm in BL. Asterisk: injected sperm, unusual for this size. (K) An individual 3.8 mm in BL carrying multiple pre-vitellogenic eggs. (L–M) Male-mature but female-immature individual. (L) Fully differentiated copulatory organ in an individual of BL 5.3 mm. (M) Sagittal section of newly and fully developed male reproductive organs. (N–O) Male- and female-mature individual. Scale bars: (A, F, G, L) 100 μm, (B, E, H, J, K, N) 200 μm, (C) 20 μm, (I) 10 μm, (L, M, O) 500 μm. Abbreviations: DS, digestive syncytium; Gd, gonoduct; Gp, gonopore; GV, granular vesicle; P, penis; Ph, pharynx; SD, sperm duct; SV, seminal vesicle; T, testes

#### Asexual Hatchlings

Hatchlings emerged from eggs at 0.5–0.7 mm in length (**Fig. 1C**). They were initially sluggish, but began gliding actively within a few hours. They exhibited key anatomical features such as an anterior condensation of neurons (e.g., Srivastava et al., 2014; Hulett et al., 2020), a pharynx occupying nearly half the body length, a statocyst, frontal gland, and digestive syncytium (**Fig. 4A**). No reproductive structures were observed at this stage. Growth rates varied among individuals, even within the same clutch. Cannibalism appeared to be an important mechanism for both accelerating growth and removing weaker individuals. When hatchlings were reared in isolation, some grew markedly slower, or even shrank and eventually disappeared, under mild stress conditions (e.g., delayed feeding or water changes) despite receiving regular food.

#### Immature Male Juveniles (1.4–3.0 mm)

Sexual development became apparent in individuals measuring 1.8–1.9 mm in BL, indicated by the appearance of testis follicles along the dorsolateral margin, just behind the anterior condensation (**Fig. 4B**). However, spermatogenesis could have started in specimens as small as 1.4 mm (**Fig. 4C**). By 2.1 mm, a rudimentary copulatory organ (CO) was present in the ventral anterior third of the body (**Fig. 4D**). This immature CO comprised two main structures connected by longitudinal muscles. The antrum was continuous with the body wall and possessed a small and slightly flared cavity. A relatively slender penis with a small, round and only slightly elongated granular vesicle which could be considered as “a young granular vesicle”, was also observed. Small pools of sperm were present posterior to the granular vesicle, but lacked a muscular wall and were hence false seminal vesicles.

#### Male-Immature, Female-Immature Juveniles (3.0–5.0 mm)

By 3.0 mm in BL, testis follicles in well-fed individuals expanded beyond the anterior dorsolateral regions and often extended across the pharyngeal region, as also observed in larger adults with multiple oocytes (**Fig. 4E**). The seminal vesicle became easily distinguishable, and the CO showed advanced differentiation (**Fig. 4F, G**). Notably, the anterior tube of the granular vesicle observed connected anteriorly to the stylet which, in turn, was connected to the gonopore by the antrum. A sclerotized stylet was also observed, although needle structure and coloration hinted at the needles, at least, being immature (**Fig. 4F**). Behavioral observations showed that no males smaller than 5.0 mm in BL exhibited any signs of copulation, so it was suggested that the male CO was not yet fully functional before reaching 5.0 mm.

The first signs of female reproductive development also emerged at 3.0 mm, indicated by the appearance of oocytes along the ventrolateral margin of the posterior third of the body (**Fig. 4H, I**). The number and size of oocytes increased with body length in this stage, but most remained pre-vitellogenic (**Fig. 4J, K**).

#### Male-Mature Young Adults (> 5.0 – 6.0 mm)

Individuals at this stage had a fully mature and functional male CO (**Fig. 4L**), and individuals began copulatory behavior (**Supplementary Movie S1**). Copulatory behavior started with exploratory probing. If the initiating worm was satisfied with the partner’s size or sexual status (although the mechanism is still unclear), a forceful copulatory stab of its penis followed. Interestingly, partners chosen by worms in this stage tended to be larger and did not reciprocate the stab. Inseminated worms may have recoiled or attempted to escape, but did not actively reject partners.

Oocyte development still continued. The number of vitellogenic oocytes was quite large in this stage. Sperm introduced through traumatic insemination, referred to as “allosperm” (Bourlat and Hejnol, 2009), was also occasionally found near some eggs (**Fig. 2G**). This observation suggests that individuals at the larger end of this developmental stage may already be receptive to copulation. However, mature eggs are absent from the medial area behind the seminal vesicle (**Fig. 4M**).

#### Mature Adults (≥ 6.0 mm)

Individuals over 6.0 mm in BL displayed fully developed male and female reproductive systems (**Fig. 4N, O**). The mature male CO was well-developed as showed in **Fig. 3**. Seminal vesicles were often large enough to be visible to the naked eye.

These individuals engaged in reciprocal mating **(Fig. 5A, Supplementary Movie S1, S2)**, forming a nose-to-tail circle, probing briefly, and then stabbing each other with stylets on the tip of male CO. Each partner usually performed only one or two copulatory attempts before disengaging. Allosperm was often detected inside the bodies of the inseminated individuals (**Fig. 5B**). Their position was clearly different from that of newly produced sperm normally stored in the reproductive tract.

**Figure 5.**
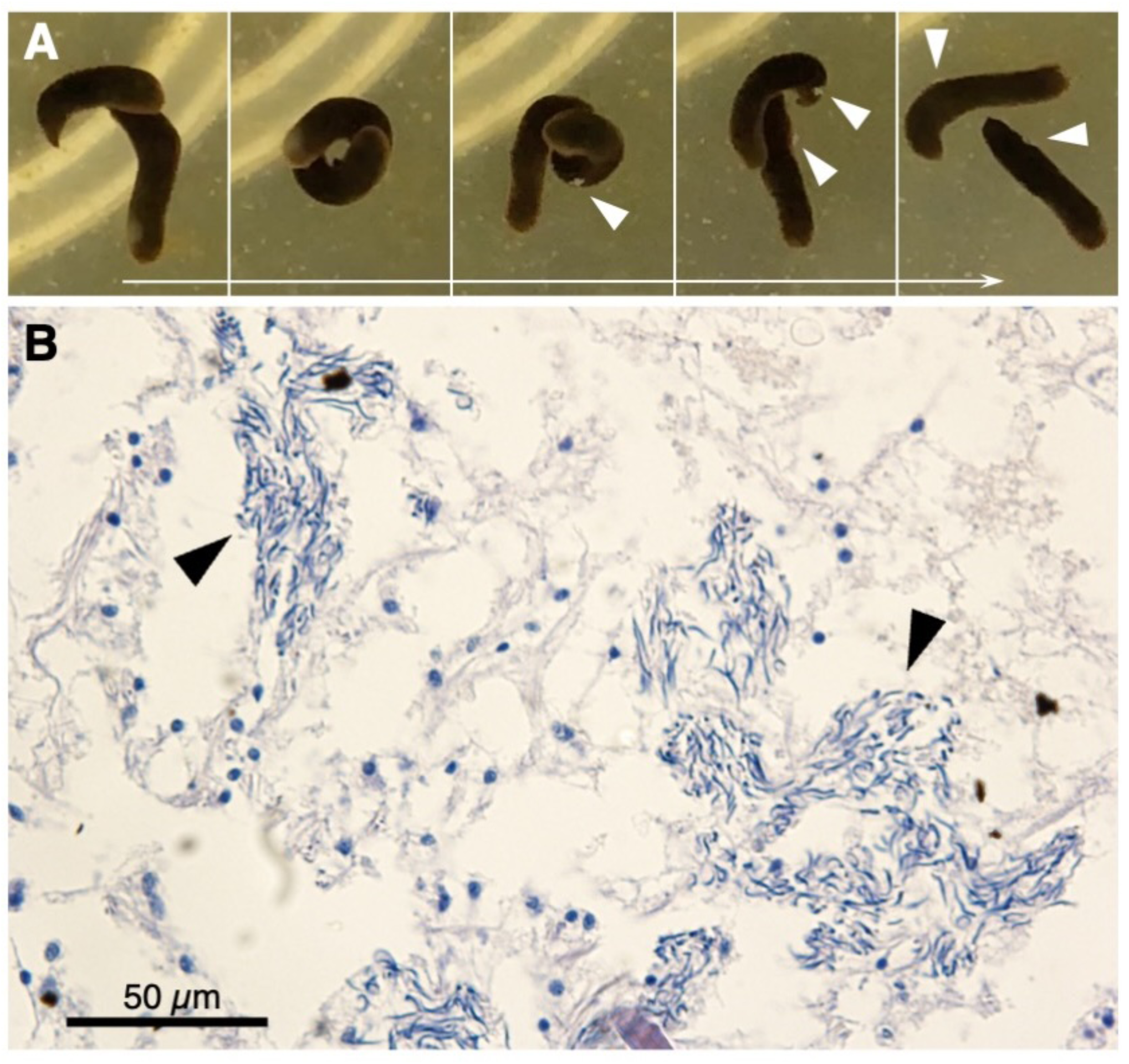
Sequences of copulatory behavior (A) and allosperm detected in an inseminated individual (B). Arrowheads in (A) indicate semen reciprocally transferred between mating partners. Arrowheads in (B) indicate sperm located within the body of the inseminated individual. The surrounding tissue is connective tissue immediately beneath the epithelium.

Individuals of this size carried multiple vitellogenic oocytes at once (**Fig. 4O**). The largest wild-collected individual was 8.0 mm, but captive worms could reach up to 14 mm and held over 20 vitellogenic oocytes. Spherical, oviposition-ready eggs with envelopes were found exclusively posterior to the male copulatory organ (**Fig. 2D**).

## Discussion

### Sexual maturation and reproductive development

In this study, we demonstrate that sexual maturation in the hermaphroditic acoel *Hofstenia atroviridis* is protandrous and depends primarily on body size rather than chronological age (**Fig. 6**). Although reproductive morphology has been described for many acoel species, developmental studies remain rare and are strongly biased toward a limited number of taxa and geographic regions, particularly within the infraorder Crucimusculata (Jondelius et al., 2011). By focusing on *H. atroviridis*, a representative of the more basal and understudied Prosopharyngida, our work helps to reduce this bias. This study represents only the third published analysis of reproductive system development in acoels and contributes to a broader understanding of reproductive evolution within Xenacoelomorpha by integrating morphological, histological, and behavioral data.

**Figure 6.**
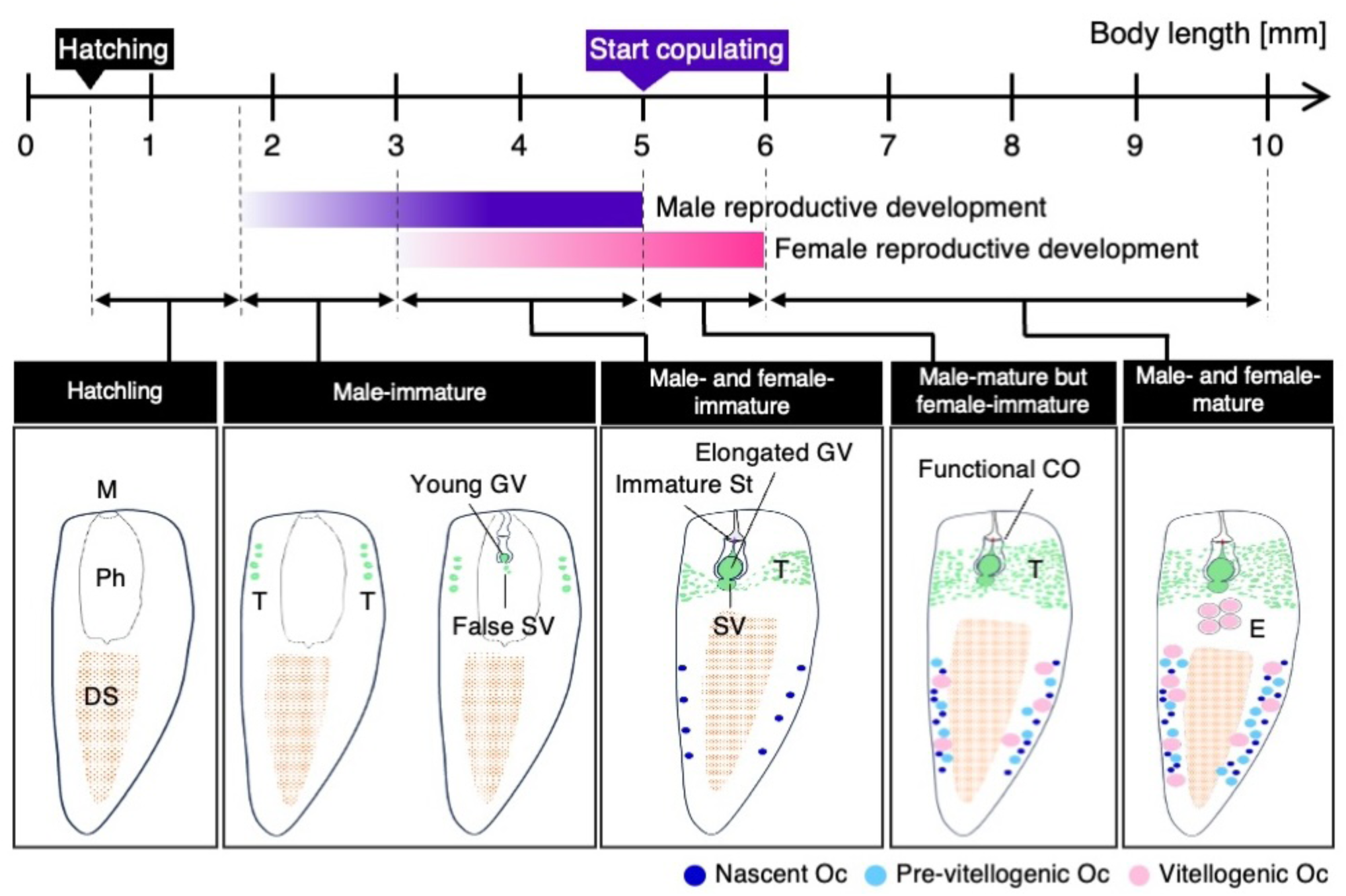
Schematic diagram of reproductive development of *Hostenia atroviridis*. Since this study revealed that the reproductive development of *H. atroviridis* is correlated to body size, diagrams of each stage are showed along with body length (BL; mm). In summary, male reproductive development starts earlier than that of female. Maturation as males is achieved and copulatory behavior is observed even in female-immature individuals (5.0-6.0 mm in BL). Individuals larger than 6.0 mm in BL have matured as both males and females. Abbreviations: DS, digestive syncytium; CO, copulatory organ; E, egg; GV, granular vesicle; M, mouth; Oc, oocyte; Ph, pharynx; St, stylet; SV, seminal vesicle; T, testis.

### Reproductive structure and cell type diversity

Like most acoels, *H. atroviridis* possesses a morphologically complex reproductive system (Atherton and Jondelius, 2022). However, our analysis reveals that this complexity is achieved with a remarkably limited diversity of cell types. The male copulatory apparatus consists almost entirely of muscle cells with a minor epithelial component, whereas follicle cells constitute the only somatic element of the female reproductive system (**Fig. 3**). Despite the absence of genital ducts, enclosed gonads, or discrete multicellular glands, this system successfully organizes germ cells, supports gametogenesis, stores gametes, and enables their transport and transfer.

This organization contrasts with that of most internally fertilizing animals, such as vertebrates, insects, and annelids, which typically possess enclosed gonads, diverse somatic support cells, specialized epithelia, and duct systems (Brusca et al., 2016; Chapman, 2013a, b; Webster and Webster, 1974). In these groups, muscles primarily serve supportive roles in gamete and secretion transport. In *H. atroviridis*, by contrast, muscles play a central role: they partially delimit gonads, move and store gametes, and constitute nearly the entire copulatory apparatus, while somatic support cells are minimal and absent from the testes. How such a simple histological organization enables efficient internal fertilization remains an important question.

### Internal fertilization without specialized ducts

Acoels are what Extavour (2007) describes as wholly internally fertilizing, i.e., gametes are deposited inside the other individual through copulation, yet defined gonoducts are absent, including in *H. atroviridis* (Achatz et al., 2012). While oocyte movement has been attributed to muscle contractions and internal pressure (Chandra et al., 2025), the transport of spermatocytes is less intuitive. Spermatocytes move as membrane-free clusters, follow a trajectory perpendicular to that of oocytes, and must converge from widely distributed testes toward a relatively small seminal vesicle.

Passive displacement by proliferating follicles is unlikely to explain this directionality. Testes develop dorso-medially, whereas the seminal vesicle is ventromedial, and follicles remain close to the body wall without approaching the pharynx (**Fig. 2**; Chandra et al., 2025). We therefore propose that parenchymal muscle fibers may link testes to the seminal vesicle, providing directional guidance for spermatocyte movement. Muscle cells in *H. miamia* express positional control genes dynamically during regeneration (Raz et al., 2017), suggesting a possible molecular mechanism for such guidance. In addition, repeated body elongation and contraction during locomotion may mechanically facilitate spermatocyte movement by altering parenchymal spacing. Together, these observations suggest that wholly internal fertilization in acoels can be achieved with reduced morphological complexity, potentially compensated by muscular and molecular guidance systems.

### Novel insights into reproductive components

Although the testis of adult *H. atroviridis* is unpaired, it originates as a paired structure during development (**Fig. 4**), as also observed in *H. miamia* (Chandra et al., 2025). Bock (1923) attributed the peripheral position of testes to an anterior shift of the copulatory organ. Our data suggest that anteriorization of both the mouth and pharynx also played an important role, with the enlarged pharynx displacing the testes laterally and causing the developmental field of the testis to split.

We further report the first instance of an acoel penis composed of three distinct muscle layers. The longitudinal outer and inner layers may contribute to sperm propulsion, whereas the thick circular layer likely generates the hydrostatic pressure required for stylet deployment. Although the innermost penial layer resembles the outer layer of body wall musculature (Ricci and Srivastava, 2021), structural differences and comparative evidence from other acoels (Chiodin et al., 2013; Mamkaev and Kostenko, 1991) suggest this similarity is superficial, consistent with the presence of a copulatory anlage in *Hofstenia*.

The granular vesicle exhibits a thin, reticulated wall labeled by phalloidin and smooth muscle actin (**Fig. 3**), consistent with an epithelial structure also reported in *H. miamia* (Chandra et al., 2025). The thick muscular wall observed in histological sections does not belong to the granular vesicle itself. Comparative data indicate that granular (prostatic) vesicles in acoels primarily serve secretory functions rather than sperm transport (Achatz et al., 2010; Hooge and Tyler, 1999; Zauchner et al., 2015), and their diversity across species suggests convergent evolution. Finally, the antrum of *H. atroviridis* is bilayered, continuous with the body wall, and lined by uninterrupted cilia (**Fig. 3**), supporting an origin through inward folding of the body wall, as proposed for acoels more broadly (Mamkaev and Kostenko, 1991).

### Seasonal variation and ecological context

Population-level studies of reproductive dynamics in acoels are lacking, limiting ecological interpretation. We observed a seasonal reduction in both size and abundance of wild-caught *H. atroviridis*. Although increased winter low-tide heights restricted access to shoreline seaweed, even diver-assisted sampling failed to recover individuals comparable in size or number to those collected in summer. This indicates genuine seasonal variation in *H. atroviridis* populations, with potential implications for growth, maturation, and reproductive timing.

### Evolution of size-based protandrous sexual hermaphroditism

Individuals of different ages but similar size exhibit comparable levels of sexual maturity (**Fig. 4, 6**), a pattern also reported for *H. miamia* (Chandra et al., 2025). Because body size in *H. atroviridis* depends strongly on environmental factors such as food availability, sexual maturation is partially uncoupled from age. Individuals can therefore reproduce as soon as they reach a sufficient size, extending the reproductive period and increasing fitness, particularly for faster-growing individuals. Moreover, size-dependent protandry allows male function to persist during periods of shrinkage, enabling reproduction in at least one sexual role under adverse conditions.

Protandry also allows energetically inexpensive sperm production to precede the energetically costly phases of growth and oogenesis. By establishing a substantial sperm reserve before oocyte maturation, energetic conflict is reduced and oocyte production can be maximized, suggesting that protandry may have evolved as a mechanism to resolve resource-allocation conflicts in the reproductive soma.

This framework alone does not explain the pronounced asymmetry between male and female reproductive morphology in *Hofstenia*. Ancestral-state reconstructions indicate that greater elaboration of male reproductive structures is ancestral in Acoela (Abalde and Jondelius, 2025). As an early-diverging lineage (Jondelius et al., 2011), *Hofstenia* likely retained this bias, which may have been further accentuated by sexual selection following the evolution of protandry (Eberhard, 1985; Leonard, 2006).

### Protandrous simultaneous hermaphroditism in Bilateria

Protandrous simultaneous hermaphroditism (PSH) is rare across Bilateria (Jarne and Auld, 2006; Sasson and Ryan, 2017), but has evolved repeatedly in diverse taxa, including shrimps, flatworms, and polychaetes (Zhang et al., 2017). Studies of these groups show that PSH is frequently associated with size-dependent, dietary, or socially mediated sex allocation (Nakashima, 1987; Bauer and Holt, 1998; Baldwin and Bauer, 2003; Zupo, 2001; Sella, 1988; Santi et al., 2018; Ramm et al., 2019). Within Acoela, protandry has been reported in multiple lineages, whereas protogyny appears rare (Kozloff, 2000). Given the large number of described species of acoels (Atherton and Jondelius, 2022), broader comparative studies will be necessary to infer general evolutionary patterns of PSH in the group.

### Implications for the urbilaterian reproductive system

While reconstructions of the ancestral bilaterian have focused largely on external morphology, far less is known about internal systems, particularly reproduction. Environmental reconstructions suggest that early bilaterians were small, motile, benthic organisms living under low-oxygen, low-food conditions (Budd and Jensen, 2015; Sperling and Stockey, 2018), a context in which hermaphroditism and internal fertilization may have been advantageous.

Recent phylogenomic evidence suggests that internally fertilizing, hermaphroditic nemertodermatids represent the earliest-diverging xenacoelomorph lineage (Redmond, 2024), strengthening support for these traits in the urbilaterian. In addition, the early emergence of a transcriptionally distinct reproductive soma (Devlin et al., 2023), together with our demonstration that complex reproductive function can arise from few cell types, raises the possibility that simple genitalia capable of internal fertilization evolved early in bilaterian history.

## Conclusions

Our findings suggest that reproductive strategies observed in extant, early-diverging bilaterians such as acoels may reflect ancient and evolutionarily stable conditions, including internal fertilization and hermaphroditism. These strategies likely provided flexibility in reproductive timing and resource allocation and may have contributed to early bilaterian diversification. Further comparative studies of reproductive development across acoels, integrated with molecular and ecological approaches, will be critical for refining hypotheses on reproductive evolution at the base of Bilateria.

## Acknowledgements

We would like to express our gratitude to H. Kohtsuka and M. Kawabata for arranging and helping our field collection. M. Nakamura, R. Kimbara, H. Yamaguchi, D. Sato and all other staffs and students of the Misaki Marine Biological Station, the University of Tokyo, supported our experimental procedures. M. Yoshida, D. Kurokawa, M. Okanishi and K. Oguchi gave us a lot of valuable comments on this study. This work was supported by Grant-in Aids for Scientific Research A (No. 18H04006) and for Challenging Research (Pioneering) (No. 21K18240) to TM from the Ministry of Education, Culture, Sports, Science and Technology of Japan.

## Competing Interests

The authors have no competing interests to declare.

## Author Contributions

RP, CH and TM designed the study and collected specimens in the field. RP and CH performed experimental procedures under the supervision of YO and TM. RP, YO and TM made figures and wrote the manuscript. All authors checked the submitted version of the manuscript, and agreed to the submission.

## Notes

### Competing Interest Statement

The authors have declared no competing interest.

