## Supplementary figures and images for "Postembryonic Development and Morphology of the Reproductive System in the Acoel *Hofstenia atroviridis*"

### Supplementary Figures 1-3

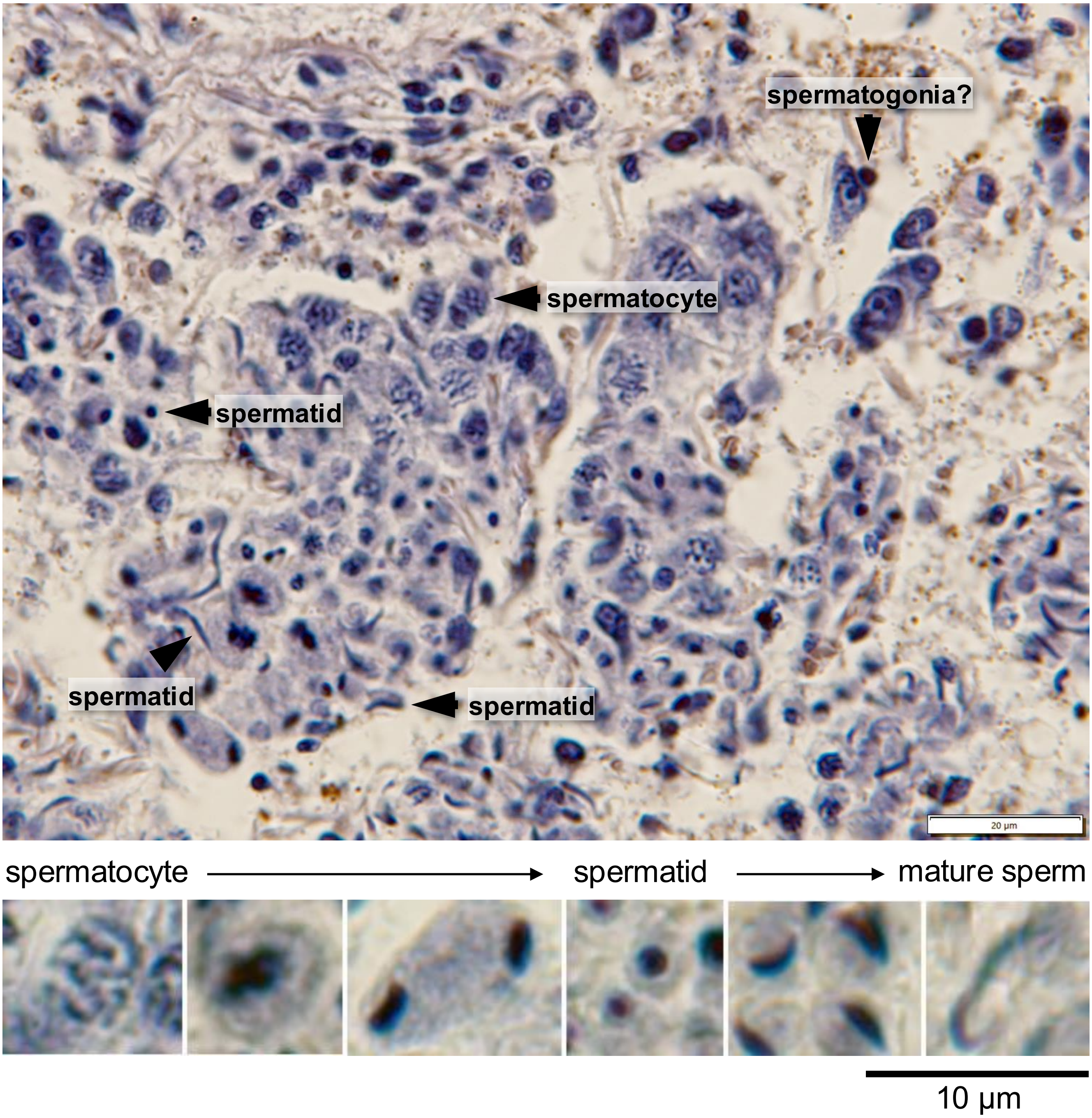


**Supplementary Figure S1**

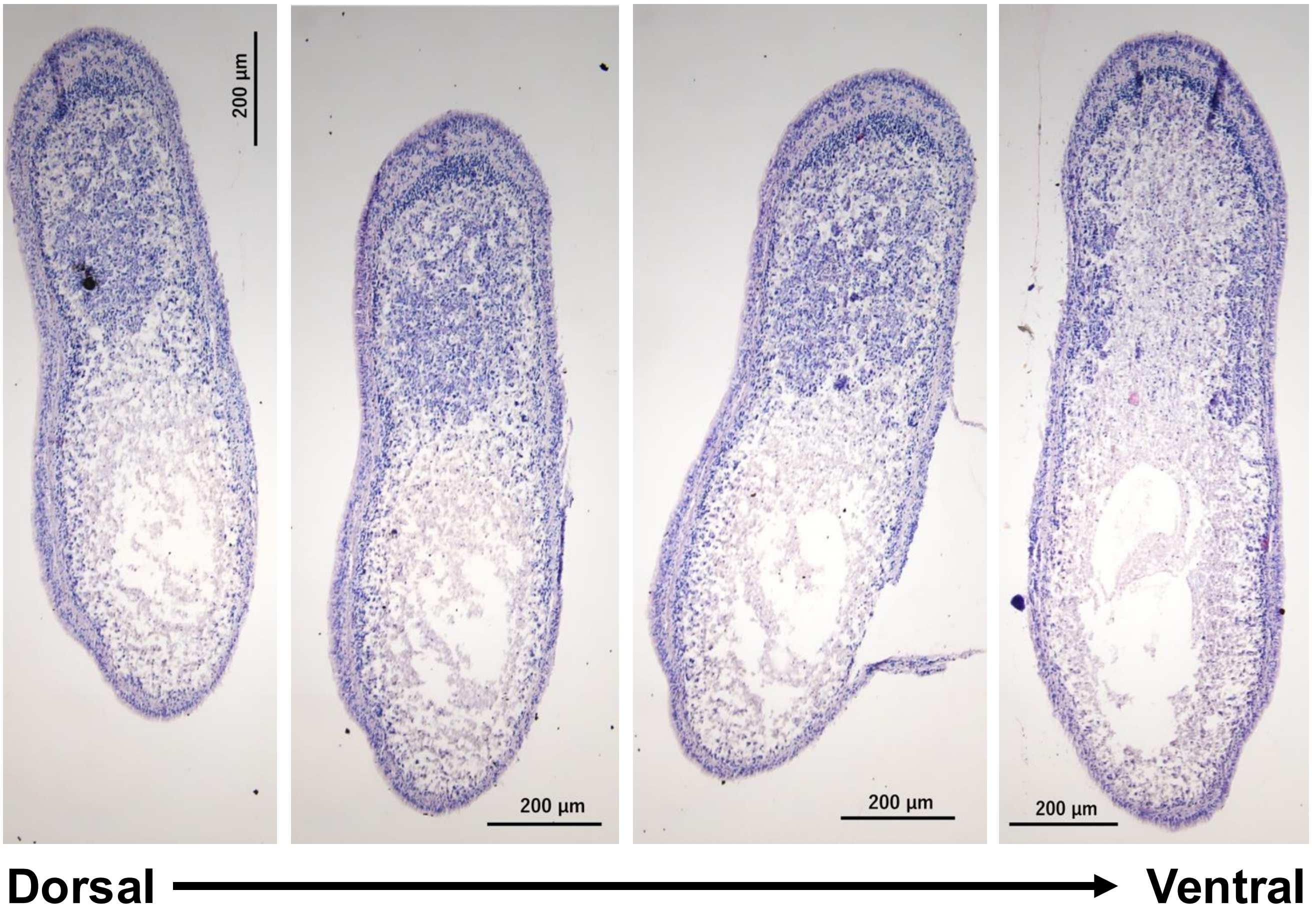


**Supplementary Figure S2**

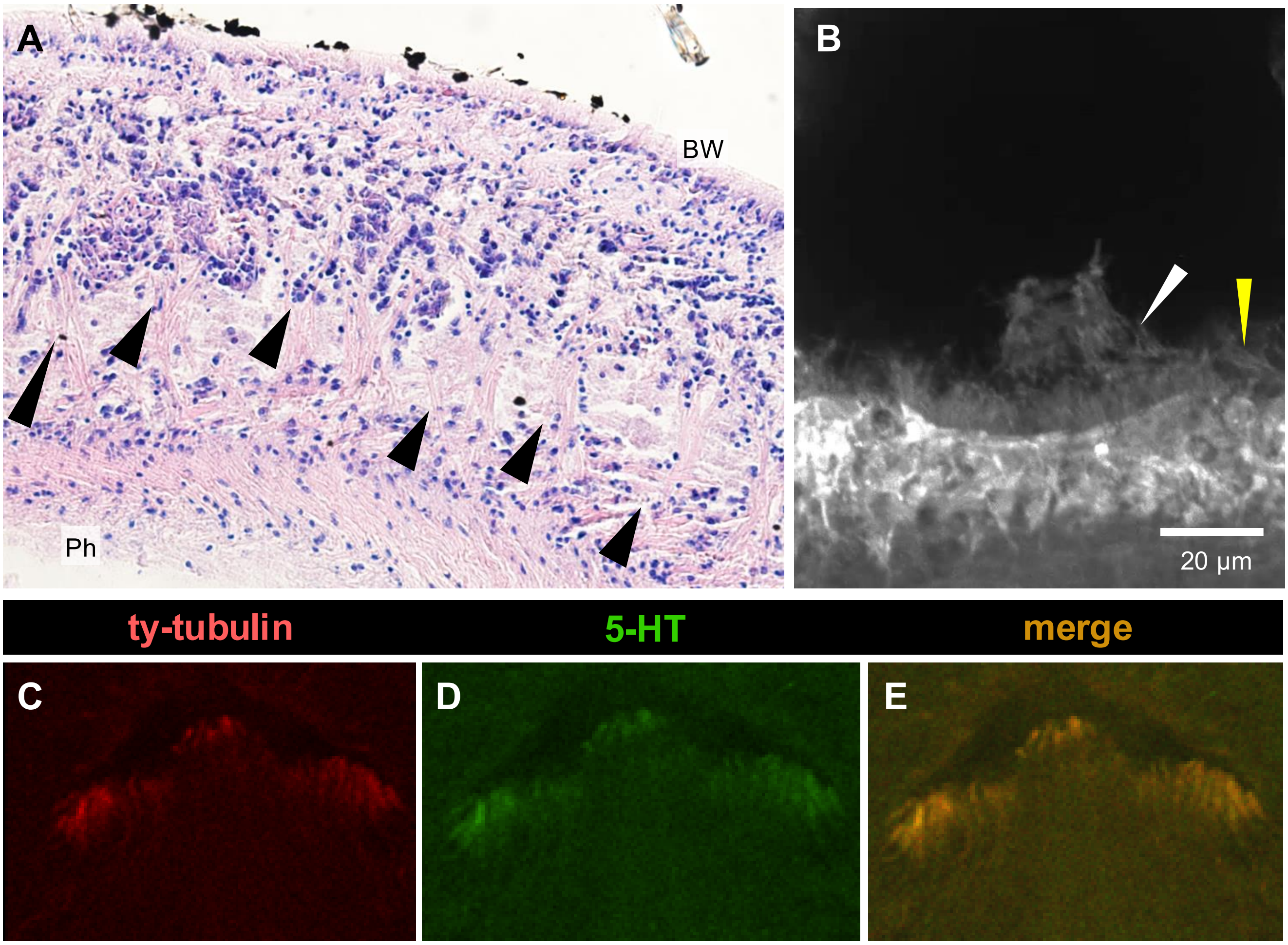


**Supplementary Figure S3**
